# DNCON2: Improved protein contact prediction using two-level deep convolutional neural networks

**DOI:** 10.1101/222893

**Authors:** Badri Adhikari, Jie Hou, Jianlin Cheng

## Abstract

**Motivation:** Significant improvements in the prediction of protein residue-residue contacts are observed in the recent years. These contacts, predicted using a variety of coevolution-based and machine learning methods, are the key contributors to the recent progress in *ab initio* protein structure prediction, as demonstrated in the recent CASP experiments. Continuing the development of new methods to reliably predict contact maps is essential to further improve *ab initio* structure prediction.

**Results:** In this paper we discuss DNCON2, an improved protein contact map predictor based on two-level deep convolutional neural networks. It consists of six convolutional neural networks – the first five predict contacts at 6, 7.5, 8, 8.5, and 10 Å distance thresholds, and the last one uses these five predictions as additional features to predict final contact maps. On the free-modeling datasets in CASP10, 11, and 12 experiments, DNCON2 achieves mean precisions of 35%, 50%, and 53.4%, respectively, higher than 30.6% by MetaPSICOV on CASP10 dataset, 34% by MetaPSICOV on CASP11 dataset, and 46.3% by Raptor-X on CASP12 dataset, when top L/5 long-range contacts are evaluated. We attribute the improved performance of DNCON2 to the inclusion of short- and medium-range contacts into training, two-level approach to prediction, use of the state-of-the-art optimization and activation functions, and a novel deep learning architecture that allows each filter in a convolutional layer to access all the input features of a protein of arbitrary length.

**Availability:** The web server of DNCON2 is at <u>http://sysbio.rnet.missouri.edu/dncon2/</u> where training and testing datasets as well as the predictions for CASP10, 11, and 12 free-modeling datasets can also be downloaded. Its source code is available at <u>https://github.com/multicom-toolbox/DNCON2/</u>.

**Supplementary information:** Supplementary data are available online.

## 1 Introduction

In recent years, protein residue-residue contacts have been identified as a key feature for accurate de novo protein structure prediction (Jones, 2001; Marks *et al.*, 2012; Michel *et al.*, 2014; Michel, Menéndez Hurtado, *et al.*, 2017; Mabrouk *et al.*, 2016). Successful de novo structure prediction methods, in the recent CASP experiments, have attributed much of their performance to the incorporation of predicted contacts (Zhang *et al.*, 2016; Xu and Zhang, 2012; Ovchinnikov *et al.*, 2016). In terms of usefulness, contacts with sequence separation of at least 24 residues, i.e. long-range contacts, have been found more useful in structure modeling and are usually the primary evaluation target for evaluating and comparing contact-prediction methods. While long-range contacts are most useful for folding proteins using fragment-based methods like FRAGFOLD (Kosciolek and Jones, 2014), Rosetta (Ovchinnikov *et al.*, 2016) and Quark (Xu and Zhang, 2012), for fragment-free methods like CONFOLD (Adhikari *et al.*, 2015) and GDFuzz3D (Pietal *et al.*, 2015), other two types of contacts - short- and medium-range contacts - are also important. While successful contact prediction methods like DNCON (Eickholt and Cheng, 2012) find it effective to predict these separately, a more recent trend of predicting all contacts with a single machine learning architecture appears promising (Jones *et al.*, 2014; Wang *et al.*, 2017).

Much of the recent improvement in the performance of contact prediction is from detecting coevolving residue pairs in a multiple sequence alignment and from the machine learning techniques used to integrate these predictions as features along with other standard features. Coevolution-based contact predictors can generally predict accurate contacts in presence of at least a few hundred effective sequences in the input alignment (Michel, Skwark, *et al.*, 2017). However, recent state-of-the-art methods demonstrate that integrating these co-evolution-based predictions with other features and using a machine learning method to make final predictions, can almost always perform better than using coevolution information alone. These integrative contact predictors have used neural networks (Jones *et al.*, 2014), random forests (Skwark *et al.*, 2014), and convolutional neural networks (Wang *et al.*, 2017) to combine co-evolutionary features with other common features like secondary structures and position specific scoring matrices.

As a successor of our deep belief network based contact predictor, DNCON (Eickholt and Cheng, 2012, 2013), which was ranked as the top method in the CASP10 experiment (Monastyrskyy *et al.*, 2014), in this paper, we present our improved contact prediction method - DNCON2. The primary enhancements of DNCON2 are (a) inclusion of coevolution-based features, (b) new deep convolutional neural networks to predict full contact maps, and (c) addition of new features at multiple distance thresholds, which further improves the performance. In DNCON2, we transform all 27 input features, e.g., scalar features like protein length, one-dimensional (1D) features like secondary structure prediction, and two-dimensional (2D) features like coevolution-based predictions, into 56 two-dimensional features. As the first step of our two-level prediction approach, we train five convolutional neural networks (CNNs), which accept these 56 2D features as input to predict contact maps at distance thresholds of 6, 7.5, 8, 8.5, and 10 Å. In the second level, a separate CNN is trained with these five sets of predictions as additional 2D features, to make final short-, medium-, and long-range predictions in one contact map all at once. Finally, we test our method using the free-modeling datasets of CASP10, 11, and 12 and compare it with other state-of-the-art methods, and, also discuss how the various training hyper-parameters influence the performance.

## 2 Methods

### 2.1 Datasets and evaluation metrics

We used the original DNCON dataset consisting of 1426 proteins having length between 30 and 300 residues curated before the CASP10 experiment to train and test DNCON2. The protein structures in the dataset were obtained from the Protein Data Bank (PDB) and of 0–2 Å resolution, which were filtered by 30% sequence identity to remove redundancy. 1230 proteins from the dataset are used for training and 196 as the validation set, and the two sets have less than 25 percent sequence identity between them. In addition to the validation dataset, we benchmarked our method using (a) 37 free-modeling domains in the CASP12 experiment, (b) 30 free-modeling domains in the CASP11 experiment (Kinch *et al.*, 2016), and (c) 15 free-modeling domains in the CASP10 experiment (Monastyrskyy *et al.*, 2014). These CASP free-modeling datasets have zero or very little identity with the training dataset.

In this study, we define a pair of residues in a protein to be in contact if their carbon beta atoms (carbon alpha for glycine), are closer than 8 Å in the native structure. We consider contacts as long-range when the paring residues are separated by at least 24 residues in the protein sequence. Similarly, medium-range contacts are pairs which have sequence separation between 12 and 23 residues and short-range contacts are pairs with sequence separation between 6 and 11 residues. These definitions are consistent with the common standards used in the field (Monastyrskyy *et al.*, 2016).

As a primary evaluation metric of contact prediction accuracy, we use the precision of top L/5 or L/2 predicted long-range contacts, where L is the length of the protein sequence. The metric has also been the main measure in the recent CASP evaluations (Monastyrskyy *et al.*, 2011, 2014, 2016) and some recent studies (Adhikari *et al.*, 2016). When evaluating the predictions for the proteins in the CASP datasets, we evaluate them at the domain level to be consistent with the past CASP assessments, although all predictions were made on the full target sequences without any knowledge of domains. We used the ConEVA tool to carry out our evaluations (Adhikari *et al.*, 2016).

### 2.2 Input features

In addition to the existing features used in the original DNCON, we used new features derived from multiple sequence alignments, coevolution-based predictions, and three-state secondary structure predictions from PSIPRED (Jones, 1999). The original DNCON feature set includes length of the protein, secondary structure and solvent accessibility predicted using the SCRATCH suite (Cheng *et al.*, 2005), position specific scoring matrix (PSSM) based features (e.g. PSSM sums and PSSM sum cosines), Atchley factors, and several pre-computed statistical potentials. During our experiments, we found PSSM and amino acid composition from the original DNCON feature set were not very useful and hence removed them from the feature list. Besides the DNCON features, the new features include coevolutionary contact probabilities/scores predicted using CCMpred (Seemayer *et al.*, 2014), FreeContact (Kaján *et al.*, 2014), PSICOV (Jones *et al.*, 2012), and alignment statistics such as number of effective sequences, Shannon entropy sum, mean contact potential, normalized mutual information, and mutual information generated using the alignment statistics tool ‘alnstat’ (Jones *et al.*, 2014). During our experiments, often, PSICOV did not converge when there are too many or too few alignments, especially if the target sequence is long. To guarantee to get some results, we set a time limit of 24 hours, and run PSICOV with three convergence parameters (‘d = 0.03’, ‘r = 0.001’, and ‘r = 0.01’) in parallel. If the first prediction (with option d = 0.03) finishes within 24 hours, we use the prediction, and if not, we use the second prediction and so on. Using all these features above as input, we predict contact maps at 6, 7.5, 8, 8.5, and 10 Å distance thresholds at first, and then use these five contact-map predictions as additional features to make a second round of prediction. Contact predictions at lower distance thresholds are relatively sparse and include only the residue pairs that are very close in the structure, whereas, contact predictions at higher distance thresholds are denser and provide more positive cases for the deep convolution neural network to learn (see **Suppl. Figure S1** for visualization of contact maps at various distance thresholds, and **Suppl. Table S2** for a full list of all the features used).

### 2.3 Generating multiple sequence alignments

Generating a diverse/informative multiple sequence alignment with a sufficient number of sequences is critical for generating quality coevolution-based features for contact prediction. On one hand, having too few sequences in the alignment, even though they may be highly diverse, can lead to low contact prediction accuracy. On the other hand, having too many sequences can slow down the process of co-evolution feature generation, creating a bottleneck for an overall structure prediction pipeline. To reliably generate multiple sequence alignments, an alignment method should produce at least some sequences in alignment whenever possible, and does not generate too many more sequences than necessary. Following a similar procedure in (Ovchinnikov *et al.*, 2016) and (Kosciolek and Jones, 2015), we first run HHblits (Remmert *et al.*, 2011) with 60% coverage thresholds, and if a certain number of alignments are not found (usually around 2L), then we run JackHMMER (Johnson *et al.*, 2010) with e-value thresholds of 1E^−20^, 1E^−10^, 1E^−4^ and 1 until we find some alignments. JackHMMER is not run if HHblits can find at least 5000 sequences. These alignments are used by the three coevolution-based methods (CCMpred, FreeContact, and PSICOV) to predict contact probabilities / scores, which are used as two-dimensional features and to generate alignment statistics related features for deep convolutional neural networks.

### 2.4 Deep convolutional neural network architecture

Convolutional neural networks (CNNs) are widely applied to recognize images with each input image translated into an input volume such that the size of the image are length and width of the volume, and the three channels (hue, saturation, and value) represent the depth. Based on such ideas, to build an input volume for each protein of any length, we translate all scalar and one-dimensional input features into two-dimensional features (channels) so that all features (including the ones already in 2D) are in two-dimension and can be viewed as separate channels. While scalar features like sequence length are duplicated to form a two-dimensional matrix (one channel), each one-dimensional feature like solvent accessibility prediction is duplicated across the row and across the column to generate two channels. The size of the channels for a protein is decided by the length of the protein. By having all features in separate input channels in the input volume, each filter in a convolutional layer convolving through the input volume, has access to all the input features, and can learn the relationships across the channels. Compared to the input volumes of images that have three channels, our input volumes have 56 channels.

We use a total of six CNNs, i.e. five in the first level to predict preliminary contact probabilities at 6, 7.5, 8, 8.5, and 10 Å distance thresholds separately by using an input volume of a protein as input, and one in the second level that take both the input volume and the 2D contact probabilities predicted in the first level to make final predictions (Figure 1(A)). Each of the six CNN networks have the same architecture, which has six hidden convolutional layers and one output layer consisting of 16 filters of 5 by 5 size and one output layer (Figure 1(B)). In the hidden layers, the batch normalization is applied, and ‘Rectified Linear Unit’ (Nair and Hinton, 2010) is used as the activation function. The last output layer consists of one 5 by 5 filter with ‘sigmoid’ as the activation function to predict final contact probabilities. Hence, our deep network can accept a protein of any length and predict a contact map of the same size. We use the Nesterov Adam (nadam) method (Sutskever *et al.*, 2013) as the optimization function to train the network.

**Figure 1.**
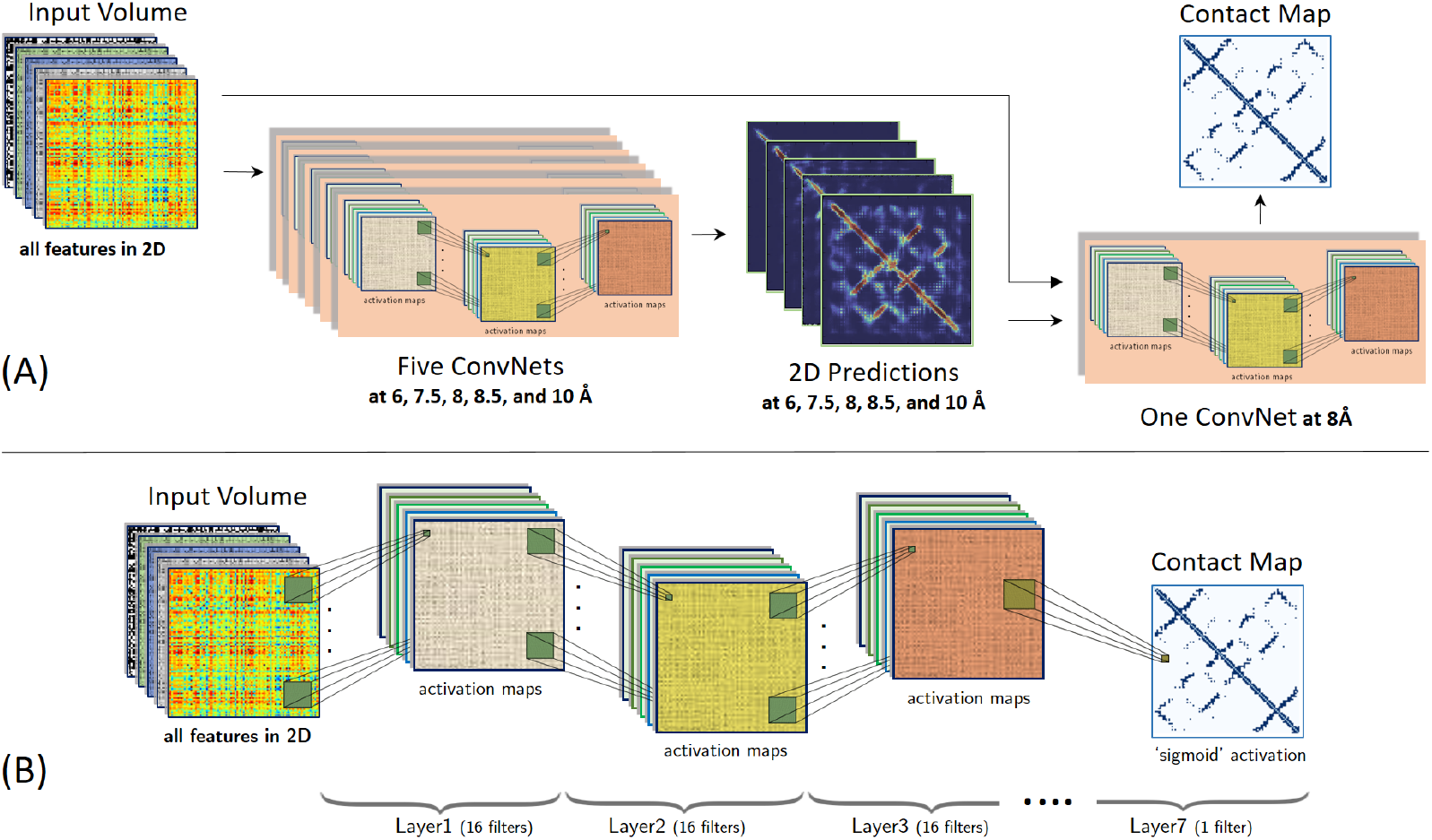
(A) The block diagram of DNCON2’s overall architecture. The 2D volumes representing a protein’s features are used by five convolution neural networks to predict preliminary contact probabilities at 6, 7.5, 8, 8.5 and 10 Å thresholds at the first level. The preliminary 2D predictions and the input volume are used by a convolutional neural network to predict final contact probability map at the second level. (B) The structure of one deep convolutional neural network in DNCON2 consisting of six hidden convolutional layers with 16 5x5 filters and an output layer consisting of one 5x5 filter to predict a contact probability map.

We train each CNN for a total of 1600 epochs with each epoch of training taking around 2 minutes. After training, we rank and select best model using the mean precision of top L/5 long-range contacts calculated on the validation dataset of 196 proteins. Our raw feature files for all 1426 training proteins use 8 GigaBytes (GB) of disk space and are expanded to around 35 GB when all features are translated to 2D format. To minimize disk input/output, we translate our scalar features and 1D features into 2D, at runtime, in CPU memory. We used the Keras library (<u>http://keras.io/</u>) along with Tensorflow (<u>www.tensorflow.org</u>) to implement our deep CNN networks. Our training was conducted on Tesla K20 Nvidia GPUs each having 5 GB of GPU memory, on which, training one model took around 12 hours. Finally, we use an ensemble of 20 trained deep models to make final predictions for testing on the CASP datasets.

## 3 Results

### 3.1 Using contact predictions at 6, 7.5, 8, 8.5, and 10 Å distance thresholds as features improves precision

A contact map is a binary version of the distance map of a protein structure according to a distance threshold, which usually is 8 Å. This threshold of 8 Å, although widely used, can be viewed as an arbitrary and rigid criterion to decide if a pair of residue is a contact or non-contact, considering the flexibility of protein structures. For instance, a pair separated by 8.1 Å distance is, by definition, a non-contact, but by 7.9 Å is a contact. And using one distance threshold causes the loss of some distance information. In order to account for uncertainty and flexibility in residue-residue distance, in a first round of prediction, using all the features and true contact maps at 6, 7.5, 8, 8.5, and 10 Å distance thresholds, we trained five CNN models to predict contact probabilities at these five distance thresholds. Then, in the second round of prediction, we added these predictions as new 2D features into the feature list and trained a sixth CNN model to predict contacts at 8 Å distance threshold.

On the 196 proteins in the validation dataset, the CNN model in the second level achieves a precision of up to 73.5 % higher than 70.7 % in the first level, when top L/5 long-range contacts are evaluated. To verify if the improvement comes from the predictions at different distance thresholds or from the iterative two-level training, in a separate experiment, we trained a second level model with only the contact prediction at 8 Å distance threshold as additional feature. In this case, a precision of 72.2 % is achieved, higher than 70.7% of using one level prediction, but lower than 73.5% of using both two-level prediction and multiple thresholds, indicating that both two-level training and multiple thresholds contribute to the improvement. The results summarized in Figure2 show similar results when top L/2 contacts are evaluated. In addition to these experiments, we tested adding more predictions at higher distance thresholds of 12, 14, 16, and 18 Å as features, and found that they did not significantly improve the performance. As an additional validation, we used the ensemble of the models trained with five distance thresholds in the first level to predict the contacts for the proteins in the validation dataset, similar to a traditional neural network ensemble in (Jones et al., 2014). Such a multi-distance ensemble has a precision of 72.8% slightly higher than the 72.6% precision achieved by an ensemble of all five models trained at the same 8 Å distance threshold, but lower than 73.5% of using predictions of multiple thresholds with two-level networks in DNCON2.

**Figure 2.**
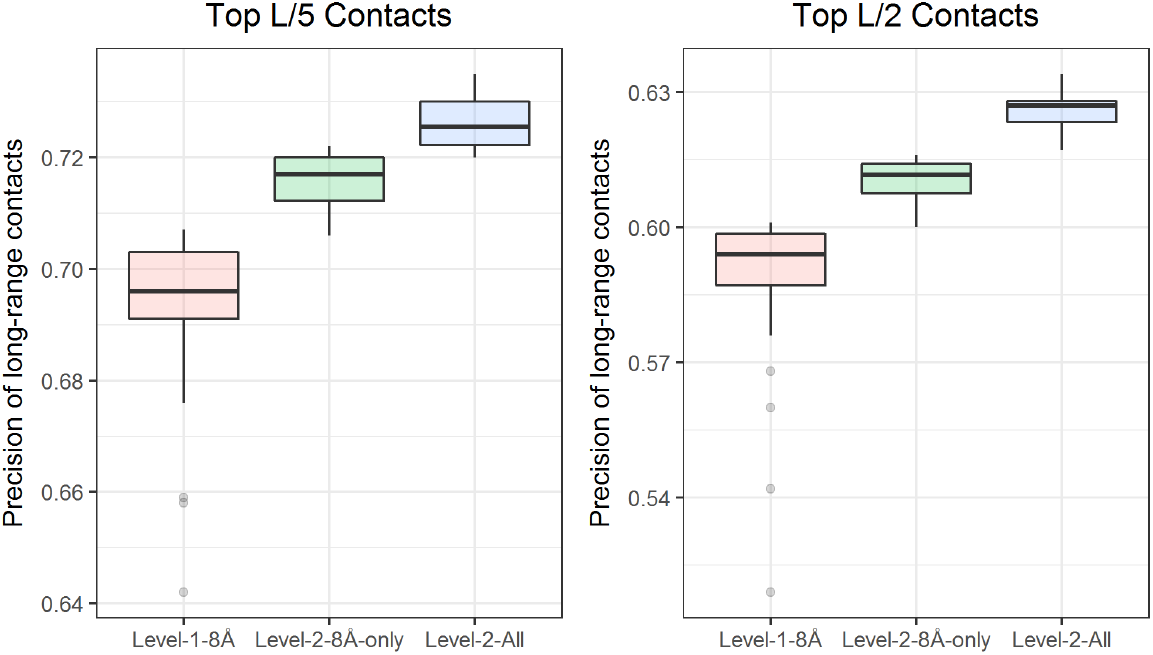
The improvement from inclusion of predictions at distance thresholds of 6, 7.5, 8, 8.5, and 10 Å as additional features, measured using the precision of top L/5 (left) and top L/2 (right) long-range contacts on the validation dataset. Box plot of precision for best 30 of 40 models for the level one model trained only using the original features (pink), the level-two model trained using only 8 Å prediction as additional feature (green), and the level-two model trained by adding all five predictions at multiple thresholds as additional features (blue).

### 3.2 Comparison between deep belief networks in DNCON 1.0 and deep convolutional neural networks in DNCON2

DNCON 1.0 used an ensemble of deep belief networks (DBN) trained with windows of seven different fixed sizes and boosting to predict contacts and achieved an accuracy of 34% on the 196 proteins in the validation dataset. For a fair comparison with DNCON, we trained one CNN using the same features that DNCON 1.0 used (excluding new coevolution-based features). Different from the DNCON 1.0 of using different networks to predict contacts at different ranges, DNCON2 uses a CNN network to predict short-, medium-, and long-range contacts of a protein of arbitrary length. With the same features as input, a CNN network trained with all contacts and non-contacts achieves a slightly better precision of 35.4 % on top L/5 long-range contacts than DNCON 1.0. So, a single CNN model performs better than a boosted and ensembled deep belief networks, suggesting that the deep convolutional neural network (CNN) is more suitable for contact prediction than the deep belief network (DBN). Moreover, it is more convenient to train and test CNN than DBN because CNN can take a full input matrix of arbitrary size as input to predict full contact maps without generating the features for each pair of residues, separating contacts at different ranges, and balancing the ratio of contacts and non-contacts as required by DBN based on fixed size windows.

### 3.3 Performance of DNCON2 on the validation and CASP datasets

On the 196 proteins in the validation dataset, DNCON2 yields a mean precision of 74%, when top L/5 long-range contacts are evaluated. As summarized in Table1, the average length, number of sequences in the alignment, and the number of effective sequences for these proteins are 190, 5,351, and 1,718 respectively. On this dataset, the three-individual coevolution-based features generated by CCMpred, FreeContact, and PSICOV can predict contacts with precisions of 51.0%, 43.1%, 42.1% respectively, for top L/5 long-range contacts, which is much lower than 74% of DNCON2. And for 96% of these proteins, DNCON2 performs better than any of the individual coevolution-based features (see **Suppl. Figure S3**). These results indicate that integrating all the 2D coevolution-based features with the other features can drastically improve the accuracy of contact prediction.

**Table 1.** Performance of DNCON2 on the 196 proteins in the validation dataset when top L/5 and top L/2 long-range contacts are evaluated. L, N, and N_eff_ stand for length of a protein, number of sequences in the alignment, and the number of effective sequences in the alignment. P_L/5_ and P_L/2_ are the precisions of top L/5 and L/2 long-range contacts.

| | L | N | $N_{\text{eff}}$ | $P_{L/5}$ | $P_{L/2}$ |
| --- | --- | --- | --- | --- | --- |
| Average | 190 | 5351 | 1718 | 74.0% | 64.2% |
| Median | 188 | 1607 | 412 | 88.2% | 74.4% |
| Maximum | 299 | 62889 | 29547 | 100.0% | 100.0% |
| Minimum | 50 | 1 | 1 | 0.0% | 0.0% |

Since predicted contacts are most useful for *ab initio* folding of proteins whose structures cannot be predicted by template-based modeling, we evaluated our method on the free-modeling protein datasets in the CASP10, 11, and 12 experiments and compared it with top CASP methods and a standard coevolution-based method MetaPSICOV (Jones *et al.*, 2014) (see Table2). Since our training and validation datasets were curated before the CASP10 experiment, the CASP datasets are independent test data. For evaluating our method on the most recent CASP12 dataset, we generated all features using all programs and databases released before the CASP12 experiment started, making our results not influenced by the new releases of protein structures and sequences thereafter. On the 37 free-modeling CASP12 domains, for which native structures were available for us to perform the evaluation, DNCON2 outperforms all the top methods participating in the CASP12 experiment such as Raptor-X (Wang *et al.*, 2017) (Table 2), MetaPSICOV (Jones *et al.*, 2014) (Table 2), iFold_1, and our own method MULTICOM-CONSTRUCT as well as the baseline method DNCON 1.0 (see **Suppl. Table S4**). Specifically, when top L/5 long-range contacts are evaluated, DNCON2 achieves an average precision of 53.4% compared to 46.3%, 42.9%, and 45.7% by Raptor-X, MetaPSICOV, and iFold_1, respectively. A similar performance is observed when top L/2 contacts are evaluated instead of L/5 (see **Suppl. Table S4**). The 24.9% precision of DNCON 1.0, which does not use any coevolution-based features, is a benchmark for the other methods, and the difference between its accuracy with the other methods highlights the improvement gained from the inclusion of the coevolution-based features into the input. The full evaluation on the complete set of CASP12 domains is presented in **Supp. Table S5**.

**Table 2.** Summary of the performance of DNCON2 on the CASP10, CASP11, and CASP12 free-modeling (FM) datasets, measured using the precision of top L/5 long-range contacts. The precision of the top method in each CASP experiment and a standard method MetaPSICOV (run locally) is also included as a reference.

| FM Dataset | Domain Count | Precision of top L/5 long-range contacts (%) |  |  |
| --- | --- | --- | --- | --- |
|  |  | Top CASP Group | MetaPSICOV | DNCON2 |
| CASP10 | 15 | 18.1 (DNCON 1.0) | 30.6 | 35.0 |
| CASP11 | 30 | 29.7 (CONSIP2) | 34.4 | 50.0 |
| CASP12 | 37 | 46.3 (Raptor-X) | 42.9 | 53.4 |

For evaluating our method on CASP11 and CASP10 free-modeling datasets, we ran MetaPSICOV locally as a benchmark. For a fair comparison, we use the same sequence databases for DNCON2 and MetaPSICOV. For completeness, we also compared DNCON2 with the best performing groups in the two CASP experiments - CONSIP2 in the CASP11 experiment (Monastyrskyy *et al.*, 2016), and DNCON 1.0 in the CASP10 experiment (Monastyrskyy *et al.*, 2014) (Table 2). On the 30 free-modeling domains in the CASP11 experiment, DNCON2 achieves an average precision of 50% compared to 34.4% by MetaPSICOV and 29.7% by the best performing method CONSIP2 (Kosciolek and Jones, 2015) in CASP11, when top L/5 long-range contacts are evaluated. Similarly, on the 15 free-modeling structural domains in the CASP10 experiment, DNCON2 achieves a mean precision of 35%, compared to 30.6% by MetaPSICOV, and 18.1% by the best performing method DNCON. For both datasets, similar results are observed when medium-range and short-range contacts are evaluated. Detailed evaluations of top L/5 and top L/2 contacts, are presented in **Suppl. Tables S6 and S7**.

### 3.4 Hyper-parameters optimization

To obtain the best performance on the validation dataset, we fine-tuned our network by investigating a range of values/options for the following hyper-parameters: (a) depth of the network, (b) filter sizes in each layer, (c) number of filters in each layer, (d) batch normalization, (e) batch size, (f) optimization function, and (g) activation function. After many rounds of iterative hyper-parameter selection, we found that the optimal parameters for number of layers was seven, filter size was five, number of filters was 16, batch size was 30, and chosen ReLU as the activation function in hidden layers, applied batch normalization at each layer, and used NAdam as the optimization function. With the performance of the CNN in a setting as a reference, we tuned each parameter, one-by-one, to study how they influenced the performance on the training data and validation data. For the depth of the network we tested networks with two to nine layers. Similarly, for filter size and number of filters, we tested filter sizes of 1, 3, 5, 7, 9 and 11, and number of filters as 1, 4, 8, 16, and 24. On the validation dataset, the networks with filter sizes greater than 3, with >=8 filters and >= 5 hidden layers deliver around the top performance (see **Suppl. Figure S8**). When the filter size is increased beyond 9, or the number of filters is increased beyond 24, or the depth of the network is increased beyond 9, the training was very slow and often failed because of insufficient GPU memory. In addition, through trials, we found that keeping the filter size in all seven layers and number of filters in the six hidden layers the same performs better than having different filter sizes or numbers of filters in different layers.

Batch normalization is important for training deep CNN to deal with the covariate shift problem (Ioffe and Szegedy, 2015). To test how batch normalization affects the training performance, we tried applying batch normalization after each layer (a), after every alternate layer (b), or only on the first layer (c), and not using batch-normalization at all (d). We found that applying batch normalization at each layer delivers the best performance compared to any of the three other settings. While the full batch normalization applied after each layer delivers a mean precision of 70.8% and the batch normalization at every alternate layer results in a mean precision of 68.7%, for top L/5 predicted long-range contacts on the validation dataset. When batch normalization is not used at all, or is applied only to the first layer, the precision drops to 65.7%. Similarly, after testing various batch sizes, we found batch sizes of around 30 delivered the best performance on the validation dataset. For optimization methods, we tested (a) ADADELTA, (b) Adagrad, (c) Adam, (d) Nesterov Adam, (e) RMSprop, and (f) stochastic gradient descent optimizers. The results show that three optimization functions Adam, Nesterov Adam, and RMSprop deliver better performance than the others, with Nesterov Adam performing best among all (see **Suppl. Figure S9**). Finally, the activation functions sigmoid, tanh, and ReLU can achieve the precisions of 70.4%, 69.4%, and 70.9% respectively, when top L/5 long-range contacts are evaluated.

Besides the machine learning hyper-parameters, we also tested if training using only long-range contacts improves the precision of long-range contact prediction. Interestingly, we find that including medium-range contacts and short-range contacts into training improves the performance even when only long-range contacts are evaluated. However, when all the local contacts are included, i.e. contacts with sequence separation less than five, we observed a slight decrease in performance (see **Suppl. Figure S10**). In summary, including all except the local contacts during training yields better precision. In addition, to test how ensembling improves the performance, we first ranked the trained models using the average precision on the validation dataset. Then, we calculated precision of the averaged predictions of top *x* models, where *x* is an integer in the range [1, 50]. The precision of ensembling increases initially and then saturates after more than four models are used.

### 3.5 Importance of features

We removed one or more features at a time and trained the CNN using the remaining features, to study the contribution of the removed features towards the overall performance of DNCON2. We tested by removing (a) multiple sequence alignment (MSA) statistics related features comprising of Shannon entropy sum, mean contact potential, normalized mutual information, and mutual information, (b) CCMpred coevolution feature, (c) FreeContact coevolution feature, (d) PSICOV coevolution feature, (e) several pre-computed statistical potentials, (f) number of sequences in the alignment and the number of effective sequences in the alignment, (g) PSIPRED and PSISOLV predictions of secondary structures and solvent accessibility, (h) PSSM related features comprising of PSSM sums and PSSM sum cosines, (i) SCRATCH secondary structure and solvent accessibility predictions, (j) relative counts of helical residues, strand residues, and buried residues, (k) sequence separation related features, and (l) length of the protein.

Our results, summarized in Figure 3, show that the features from multiple sequence alignment related statistics together are more important than any single coevolution-based features (CCMpred, FreeContact, or PSICOV) and other features. But if all three coevolution-based predictions (CCMpred, FreeContact, and PSICOV) are removed at the same time (not shown in the figure), the precision drops most (i.e., from 60% to 38%), when top L/2 long-range contacts were evaluated, suggesting the co-evolution features as a whole have the biggest impact. Among the three coevolution-based features CCMpred, FreeContact, and PSICOV, the first two (CCMpred and FreeContact) contribute equally to the overall performance. We find the length feature is unimportant. Sequence separation related features and relative counts of helical, strand, and buried residues, also do not contribute much to the performance either. Secondary structure predictions from both methods SCARTCH and PSIPRED are useful, and complement each other to improve the overall performance.

**Figure 3.**
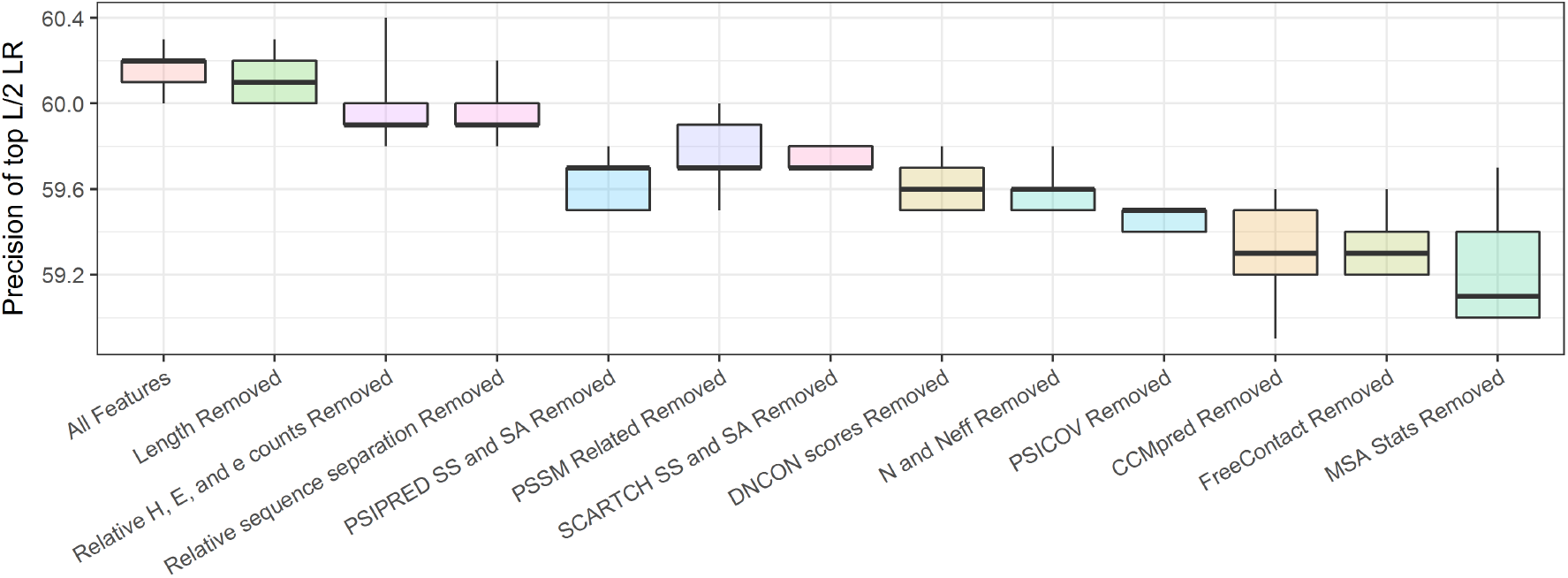
Importance of features measured by the best of five precisions of top L/2 long-range contacts on the validation dataset after removing a feature or a set of features. ‘MSA Stats’ features are multiple sequence alignment (MSA) statistics related features comprising of Shannon entropy sum, mean contact potential, normalized mutual information, and mutual information, ‘DNCON scores’ are set of several pre-computed statistical potentials, N and Neff are number of sequences and effective number of sequences.

## Conclusion

We developed DNCON2 - a new two-level deep convolutional neural network method - to predict the contact map of a protein of any length by integrating both residue-residue coevolution features and other features such as secondary structures, solvent accessibility, and pairwise contact potentials. The method can predict all the contacts in a protein at once from the entire input information of a protein, which is more effective and easier to train and use than local fixed window-based approaches such as deep belief networks. By including new coevolution features, using CNNs of multiple-distance thresholds, integrating all the features of all the residue pairs of a protein through 2D-convolution in a two-level architecture, and adopting the latest optimization and training techniques, DNCON2’s accuracy is more than double of that of DNCON 1.0 on the same validation dataset. On the three independent CASP datasets, DNCON2 outperforms the top methods in CASP10, CASP11, and CASP12 experiments. The results demonstrate DNCON2 and its deep convolutional neural network architecture are useful for protein contact prediction.

## Acknowledgements

Some of the computation for this work was performed on the high-performance computing infrastructure provided by Research Computing Support Services and in part by the National Science Foundation under grant number CNS-1429294 at the University of Missouri, Columbia MO). We would like to thank the RCSS team for their infrastructure and technical support.

